# Holotomography Reveals Protein Concentration Changes in Cardiac Spheroids under Ischemic Stress

**DOI:** 10.64898/2026.06.09.731124

**Authors:** Kenza Sackho, Paola Campagnolo, Youngchan Kim

## Abstract

Multicellular spheroids better recapitulate native cardiac tissue than two-dimensional systems, preserving cell–cell and cell–matrix interactions and relevant signalling. However, analytical tools for extracting quantitative data from these complex models remain limited. Here, we present an optimised holotomography (HT) workflow for fixed spheroids, enabling label-free quantification of protein concentration and dry mass across conditions. Using a hypoxia-reperfusion injury model to mimic myocardial infarction, HT measurements reveal a statistically significant reduction in the protein concentration of cardioids, reflecting impaired structural integrity and declining viability, subtle changes often missed by conventional approaches. These findings establish HT as a robust, scalable method for quantitative analysis of 3D cardiac models, with direct relevance for disease modelling and preclinical research.

## 1. Introduction

Two-dimensional (2D) cell cultures have long been employed in cardiovascular research, providing essential insights into cellular processes and disease progression. However, therapeutic advances in cardiovascular medicine have stagnated in recent decades, partly due to the limited translational fidelity of preclinical findings from two-dimensional (2D) systems^1^. The complex three-dimensional (3D) architecture and cellular diversity of cardiac tissue cannot be adequately recapitulated in monolayer culture. Consequently, 2D models inherently lack physiologically relevant extracellular matrix (ECM) – cell interactions, accurate toxicological responses, and the anatomical organisation of cells observed *in vivo*^2,3^.

In contrast, spheroids and organoids are 3D multicellular aggregates that more closely resemble the structural and functional microenvironment of native cardiac tissue^3^. These systems maintain cellular and molecular characteristics more faithfully than traditional 2D cultures and have become valuable tools for modelling physiology, pathology, and drug response *in vitro*^4–7^. In cardiovascular research, cardiac spheroids incorporating cardiomyocytes, endothelial cells, and fibroblasts have been developed and characterised ^8–11^. These cardiac spheroids have demonstrated microvascular formation, functional cardiomyocyte alignment, appropriate ECM deposition and toxicological responses presenting a promising platform for therapeutic and toxicological evaluation^3,8–10,12–14^. To further improve translational relevance, epicardioids, spheroids incorporating an additional epicardial layer, have since been established^10,12,13,15^. The epicardium plays critical roles in cardiac development and repair, and epicardial inclusion enables more accurate modelling of these processes. Immunofluorescence analyses using epicardial markers WT1 and TBX18 have shown that epicardial cells self-organise at the spheroid periphery and undergo epithelial-to-mesenchymal transition (EMT), mirroring native developmental programs^13,15^.

Imaging is indispensable for characterising spheroids and assessing their organisation before and after physiological or experimental perturbations. Traditionally, fluorescence-based imaging remains the method of choice owing to its molecular specificity and capability for 3D morphological reconstruction. Nonetheless, its utility is often hampered by photobleaching and phototoxicity^16,17^. To overcome these challenges, label-free optical imaging modalities, such as holotomography (HT), have emerged as powerful alternative solutions^18^. By exploiting the intrinsic refractive index properties of biological materials, these techniques also provide quantitative visualisation of live specimens without the need for exogenous contrast agents^19^.

Specifically, HT represents a 3D extension of quantitative phase imaging (QPI), a label-free optical modality that enables quantification of biomass distribution within biological specimens^20^. By quantifying the phase shift of light transmitted through a sample, HT generates spatially resolved maps of dry mass and refractive index, offering unique insights into the biochemical components of samples^21,22^. The optical phase delay retrieved from HT is proportional to the dry mass of the sample. Cell mass and density reflect cytoskeletal integrity and metabolic status, these parameters serve as sensitive biophysical indicators of cellular health^20,22,23^. As a result, changes in biomass distribution can therefore be leveraged to infer dynamic processes, including cell-cycle progression, apoptosis, and invasion behaviour. Notably, while HT has been successfully applied to visualise lung and intestinal organoids at single-cell resolution over extended time periods^24,25^, its application for quantifying biomass changes in cardiac spheroids, particularly under pathological conditions, remains largely unexplored.

In this study, we optimise an HT-based workflow for fixed organoid and spheroid samples to extract quantitative protein concentrations under both control and disease-mimicking conditions. Utilising a hypoxia-reperfusion injury model, we show that ischemic stress induces a statistically significant, measurable decrease in protein concentration, reflecting compromised structural integrity and cellular viability. Together, these findings demonstrate HT as a sensitive, intrinsically quantitative imaging modality for the rigorous analysis of 3D multicellular spheroids, offering a valuable tool for disease modelling and drug discovery.

## 2. Materials & Methods

### 2.1 Cell Culture

Cell lines were maintained at 37 °C in a humidified incubator with 5% CO_2_. Upon thawing, cells were transferred to a 15 mL conical tube containing 9 mL of pre-warmed growth medium and centrifuged at 300 × *g* for 5 min. The cell pellet was then resuspended in fresh medium before seeding. For passaging, cells were washed with phosphate buffered saline (PBS, VWR), incubated with trypsin (ThermoFisher) for approximately 3 min until detachment, and neutralised with an equal volume of growth medium before centrifugation at 300 × *g* for 5 min prior to reseeding.

Neonatal human dermal fibroblasts (NDHFs,Promocell) were cultured in Fibroblast Growth Medium 2 (FGM-2, PromoCell) and maintained for up to 9 passages, all experiments were conducted between passages 6–9. Human umbilical vein endothelial cells (HUVECs, Promocell) were cultured in Endothelial Cell Growth Medium 2 (EGM-2, PromoCell) on T75 flasks and used between passages 5– 9. H9C2 cardiomyoblasts (ATCC) were cultured in Dulbecco’s Modified Eagle’s Medium High Glucose (DMEM-HG, Merck) supplemented with 10% heat-inactivated fetal bovine serum (FBS) and 1% L– glutamine–Penicillin–Streptomycin (Merck) and used between passages 27–31.

### 2.2 EPDC Isolation

Epicardial-derived cells (EPDCs) were isolated from hearts of 4–6-week-old pigs of both sexes, supplied post mortem by The Pirbright Institute (Pirbright, UK). The collection of these tissues is controlled by the Animal Welfare and Ethical Review Board (AWERB) of The Pirbright Institute and pigs are humanely killed under the auspices of Establishment License X24684464. Hearts were transported in ice-cold transport medium (DMEM-HG (Merck) supplemented with 2% penicillin–streptomycin (P/S, Merck)). Tissue was transferred in laminar flow hood and washed in transport medium prior to dissection. The epicardial layer was carefully removed from both ventricles and auricles using forceps and transferred to a drop of PBS in petri dishes. Epicardial tissue was minced into 4-6 mm^2^ fragments using a sterile scalpel and 12-14 epicardial tissue fragments were plated onto 0.1% gelatin-coated petri dishes (Merck). Tissue fragments were allowed to adhere for 4 h to the surface prior to the addition of EPDC culture medium consisting of a 1:1 mixture of DMEM low glucose (DMEM-LG, Merck) and Medium 199 (M199, Merck), supplemented with 1% P/S, and 10% heat-inactivated fetal bovine serum (FBS).

After 3 days, the medium was replaced with EPDC culture medium containing 20% FBS. Epicardial outgrowth was monitored until confluence. All cells were detached using TrypLE Express (Gibco), neutralised with EPDC medium, and centrifuged at 300 × *g* for 5 min. EPDCs were used at passage 1.

### 2.3 Spheroid Formation Techniques

Two spheroid models were generated: Cardioids and EpiCardioids. Cardioids consisted of 12,000 H9C2 cells, 12,000 HUVECs, and 12,000 NDHFs. EpiCardioids were formed by supplementing the Cardioid base with 1,000 porcine EPDCs. All spheroids regardless of technique were cultured in DMEM-HG:M199 (4:1, v/v) supplemented with 1% insulin–transferrin–selenium (ITS, Fisher Scientific) and 1% L–glutamine–Penicillin–Streptomycin (Merck), referred to as cardiomyocyte media. For initial brightfield comparison images were taken 24 h after formation. Spheroid taken for holotomography assessment were fixed in Intracellular (IC) fixation buffer (Invitrogen) for 60 min.

#### 2.3.1 Pellet Method

Cell suspensions (500 μL per spheroid) in cardiomyocyte medium were dispensed into 1.5 mL standard microcentrifuge tubes and centrifuged at 300 × *g* for 3 min at room temperature (RT) with lids loosened. Tubes were then incubated to permit spheroid formation. After 24 h, the resulting spheroids were transferred to low-attachment, flat-bottom 96-well plates.

#### 2.3.2 Hanging-Drop Method

Cell suspensions for individual spheroids were prepared in cardiomyocyte medium and dispensed in 10 μL drops of cardiomyocyte media onto the inverted lid of a 60 mm petri dish. The base of the dish was filled with 5 mL PBS to maintain humidity. The lid was then placed onto of the dish and incubated for 24 h under standard conditions. Then, the spheroids that formed on the lid were collected and transferred to low-attachment wells containing 100 μL of cardiomyocytes medium for imaging.

#### 2.3.3 Magnetic Levitation

Once all cell lines reached confluent, NanoShuttle (Greiner) was added at 1 μL per 10,000 cells, and the cells were centrifuged in a mini centrifuge min at 100 x *g* for 5 min. The cells were then resuspended then centrifuged again, with this washing step repeated three times in total. In a 1.5 mL standard microcentrifuge tube, the different cell lines were combined in cardiomyocyte media at the required ratio, and 100 μL of this suspension was plated into low-attachment well plates (Greiner), a further 150 μL of cell medium was then added into each well. A magnet array plate holder (Greiner) was applied to levitate the cells and promote spheroids formation. Spheroids formed after 24 h.

#### 2.3.4 Ultra-Low Attachment

Confluent cells were detached and combined at the appropriate ratios in 100 μL of cardiomyocyte medium, then pipetted directly into the centre of ultra-low attachment (ULA) U-bottom 96-well plates (faCellitate). Spheroids formed within 24 h and were imaged by brightfield microscopy for comparison across methods. Following spheroid method optimisation, the ULA method was selected as the optimal model for further experimentation. To investigate the effect of spheroid size, a reduced-size condition was established in which the same cell ratios were maintained at one-quarter of the original seeding density. These smaller spheroids were cultured for 24 h prior to fixation, optical clearing, and HT assessment.

### 2.4 Viability

Spheroid viability was assessed using Calcein AM (Invitrogen) at a final concentration of 10 µM. Spheroids were incubated for 40 min with the staining solution diluted in culture medium in an incubator at 37 °C. Staining solution was then removed and spheroids were washed twice with culture medium, and incubated for a further 40 min in the same culture medium in an incubator at 37 °C. Spheroids were then imaged on a Nikon Eclipse TS2 microscope.

Cell death within spheroids was assessed using ethidium homodimer-1 at 1 µM using the same method outlined above.

### 2.5 Spheroid Brightfield Measurements

All brightfield images were taken on a Nikon Eclipse TS2 microscope and analysed using ImageJ software. Images were converted to threshold binary images, and spheroid area and roundness were then measured using the selection tool.

### 2.6 Hypoxia-reperfusion

The ULA method was selected as optimal and was therefore used to model ischemic stress. Spheroids were first cultured under normoxic conditions for 3 days, then transferred to a hypoxic incubator (<1% O_2_) for 16 h, followed by 24 h of reperfusion under normoxic (21% O_2_) conditions. Spheroids were subsequently fixed and sectioned in accordance with the protocol described below. Control spheroids were maintained under normoxic conditions throughout the culture period.

### 2.7 Whole Spheroid Staining Protocol

Spheroids were transferred to microcentrifuge tubes and fixed in IC Fixation Buffer (Invitrogen) for 60 min. Fixation was followed by an overnight blocking-permeabilisation step at 4 °C on a shaker in washing buffer (WB) consisting of 10% goat serum (Merck), 0.1% Triton X-100 (Merck), and 0.05% Tween (Merck). Spheroids were then washed three times in PBS, stained with 2 µg/mL DAPI (Merck) in PBS for 6 h, and washed three times for 5 min each in PBS. Gene Frames (Fisher Scientific) were placed between the slide and the coverslip to preserve the 3D structure of the organoids. Samples were mounted in 20 μL of mounting medium (Invitrogen) unless otherwise stated for the clearing optimisation experiments.

### 2.8 Spheroids Clearing

For the clearing comparison experiments the spheroid staining protocol was slightly adjusted to either use a ScaleS clearing protocol or a Glycerol-fructose protocol.

#### 2.8.1 ScaleS

Following fixation, spheroids were placed in S0 solution overnight. Permeabilisation was performed sequentially: samples were immersed in A2 solution for 24 h at 37 °C, followed by incubation in B4 solution for 24 h at 37 °C, and finally in A2 solution overnight at 37 °C. Samples were then washed in PBS for 6 h at room temperature (RT) and blocked overnight at 37 °C in ScaleS blocking solution. Spheroid were then placed in 2 µg/mL DAPI diluted in ScaleS blocking solution overnight at 37 °C. Samples were washed in fresh AbScale for 6 h at RT, re-blocked twice (2 h each) in ScaleS blocking solution, and refixed in 4% PFA for 1 h at RT, followed by an overnight PBS wash at 4°C. For refractive index (RI) matching, spheroids were incubated overnight at RT in ScaleS S4 solution and mounted onto glass slides. All steps were performed under gentle agitation to ensure uniform reagent penetration. Mounted spheroids were allowed to equilibrate to RT overnight. The solutions used for the ScaleS clearing protocol are listed in Table 1.

**Table 1.** Solution for ScaleS clearing.

| Solution | Reagents | pH |
| --- | --- | --- |
| S0 | 20% (w/v) D-sorbitol (Merck), 5% (w/v) glycerol (Merck), 3% (v/v) DMSO in PBS | 7.2 |
| A2 | 10% (w/v) glycerol, 4 M urea (Merck), 0.1% (w/v) Triton (Merck) X-100 in ddH<sub>2</sub>O | 7.7 |
| B4 | 8 M urea in ddH<sub>2</sub>O | 8.4 |
| ScaleS blocking solution | 2.5% (w/v) goat serum, 0.05% (w/v) Tween 20, 0.1% (w/v) Triton X-100 in PBS | 7.4 |
| AbScale | 0.33 M urea, 0.1% (w/v) Triton X-100 in PBS | 7.4 |
| <u>ScaleS S4</u> | 40% (w/v) D-sorbitol, 10% (w/v) glycerol, 4 M urea, 15% DMSO in ddH<sub>2</sub>O | 8.1 |

#### 2.8.2 Glycerol Fructose

Spheroids were stained using the spheroid staining protocol (section 2.7) then mounted in glycerol fructose clearing solution (60% (v/v) glycerol; Merck and 2.5 M fructose; Merck) for minimum 20 min at RT.

### 2.9 Analysis of Clearing Method

Spheroids were stained with DAPI, and clearing protocols were compared against control spheroids that had undergone standard permeabilisation only. Analysis was performed in the open-source image processing software ImageJ by measuring the mean gray values (MGV) of the DAPI stained channel on maximum-intensity projection, with each spheroid outlined manually. Maximum-intensity projections were then generated across defined depth ranges of the spheroid, and the MGV was measured using the same approach.

### 2.10 Preparation of Spheroid Slices

Fixed spheroids were cryoprotected in 30% sucrose (Sigma) until they sank, embedded in optimal cutting temperature compound, frozen in precooled 2-methylbutane, and stored at −80 °C. Spheroids were subsequently sectioned at 30 µm on a cryostat, and the section were collected onto slides and stored at −20 °C until further use. Prior to HT imaging, slides were equilibrated to RT for 20 min and then mounted in 20 μL of mounting medium.

### 2.11 HT Data Acquisition and Image Reconstruction

3D refractive index tomograms were obtained with a low-coherence HT system (HT-X1, Tomocube Inc.) capable of imaging thick specimens with reduced speckle noise. HT-X1 employs a 450-nm LED illumination module that is integrated with a digital micromirror device (DMD) at the pupil plane and a custom-designed condenser lens (NA = 0.72, working distance = 30 mm). By combining this illumination module with a motorised objective (UPLXAPO40X, Olympus), HT-X1 is used to acquire multiple intensity stacks of thick specimens corresponding to uniquely modulated illuminations. Each modulation is optimised to encode specific angular information of the specimen. In this work, we imaged the specimens on glass slides with 1.5# coverslips (Tomodish, Tomocube Inc.). All motorised microscopic operations were digitally controlled and monitored by an operating software TomoStudio X (Tomocube Inc.). Spheroids were located manually with the help of a brightfield preview image scanned over the area of 4 mm × 4 mm. Randomly selected organoids within the preview were imaged with a field of view (FOV) of 165 μm ×165 μm and a depth range of 140 μm. As most spheroids did not fit within a single FOV, multiple 3D data were acquired to construct a stitched image. The stitching process was automatically done by the software TomoStudio X. The system takes advantage of partially coherent illumination to encode information corresponding to multiple illumination directions simultaneously. This ensures the diversity and redundancy of information needed for accurate 3D refractive index reconstruction. An integrated inverse solver decodes the 3D refractive index information from the intensity stacks according to the Fourier diffraction theorem.

### 2.12 Pierce^™^ BCA Protein Assay

For each experiment, 8 spheroids were pooled together in a 1.5 mL microcentrifuge tube and lysed in RIPA buffer. Total protein concentration was quantified using the Pierce™ BCA Protein Assay Kit (Thermo Fisher) in accordance with the manufacturer’s protocol.

### 2.13 Data and Statistical Analysis

Segmentation and quantification of dry mass, refractive index, and protein concentration were performed using TomoAnalysis (Tomocube Inc.). Unless otherwise stated, all data are presented as individual spheroid values with the mean ± standard deviation (SD), where *n* denotes the number of individual spheroids assessed per condition, drawn from a minimum of three independent biological replicates. Data were assessed for normality using the Shapiro-Wilk test prior to statistical analysis. Statistical significance more than two groups was determined by one-way or a two-way ANOVA with Šídák’s post-hoc correction for multiple comparisons, while comparison between two groups were performed using unpaired, two-tailed Students’ t-test. In some instances, unequal sample sizes reflect sample loss during preparation, and do not result from selective exclusion. A *p*-value of < 0.05 was considered statistically significant (\**p* ≤ 0.05, \*\**p* ≤ 0.01, \*\*\**p* ≤ 0.001, \*\*\*\**p* ≤ 0.0001). All statistical analyses and graphical representations were performed using GraphPad Prism v10.3.1 (GraphPad Software Inc.). Specific statistical details for each experiment are provided in the respective figure legends.

## 3. Results

### 3.1 U Bottom and NanoShuttle Spheroids Produced Uniform Spheroids

To identify the most suitable method for generating spheroids compatible with holotomography (HT), four protocols were evaluated: pellet, hanging-drop, ultra-low attachment (ULA), and magnetic levitation (Table 2). Cardioids comprising H9C2 cardiomyoblasts, HUVECs, and NDHFs were used throughout this optimisation stage.

**Table 2.**
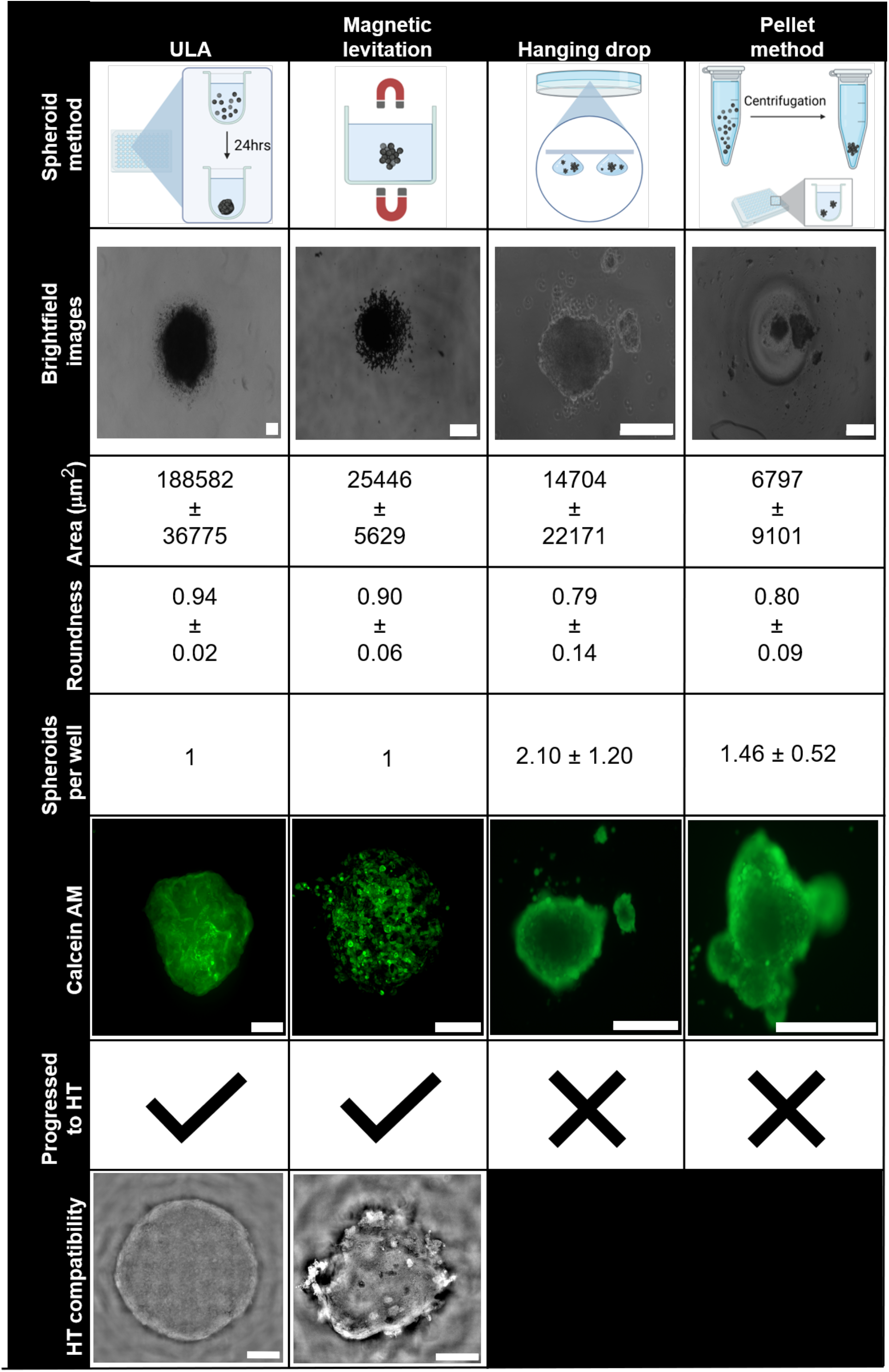
Comparison of spheroid formation techniques. Brightfield images, quantitative measurements of area, circularity, and roundness, Spheroids per well, Calcein AM viability staining, and HT compatibility are presented for each method. Scale bars: 100 µm. ULA, ultra-low attachment, HT, Holotomography

Both the hanging-drop and pellet methods produced viable spheroids, as confirmed by Calcein AM staining; however, neither consistently yielded single, discrete spheroids per well. Both methods showed marked variability in spheroid number and morphology across replicates, reflected by the high variance in area (table 1). These methods were therefore not pursued further. By contrast, both magnetic levitation and ULA plates reliably produced single, viable spheroids per well with greater morphological consistency (table 1). HT imaging revealed that spheroids generated by both methods exhibited substantial optical scattering, which attenuated light penetration and limited the quality of refractive index reconstructions. ULA-derived spheroids demonstrated modestly reduced scattering relative to magnetic levitation counterparts. To address this, subsequent experiments investigated the combined effect of reduced spheroid size and optical clearing on light penetration and reconstruction quality.

### 3.2 Optical Clearing is Incompatible with HT Analysis

To further reduce the scattering seen in ULA spheroids, small spheroids were generated in ULA plates using one-quarter of the cell density employed for the spheroids described above. In parallel, spheroids were treated with two RI-matching clearing solutions: glycerol–fructose and ScaleS. Confocal maximum-intensity projections were first acquired to assess the effect of each protocol on fluorescence signal integrity. Scales exhibited a trend toward improved fluorescent signal, although this did not reach statistical significance, whereas glycerol–fructose clearing produced a statistically significant increase in DAPI fluorescence intensity relative to uncleared controls (Fig. 1A-C). Depth-resolved analysis across the spheroid z-plane revealed that glycerol–fructose enhanced fluorescence intensity throughout 20–80% of spheroid depth, whereas the improvement with ScaleS was confined largely to the 40–80% range. Given the superior performance of the glycerol–fructose buffer in this preliminary study, glycerol–fructose was selected for HT experiments.

**Figure 1.**
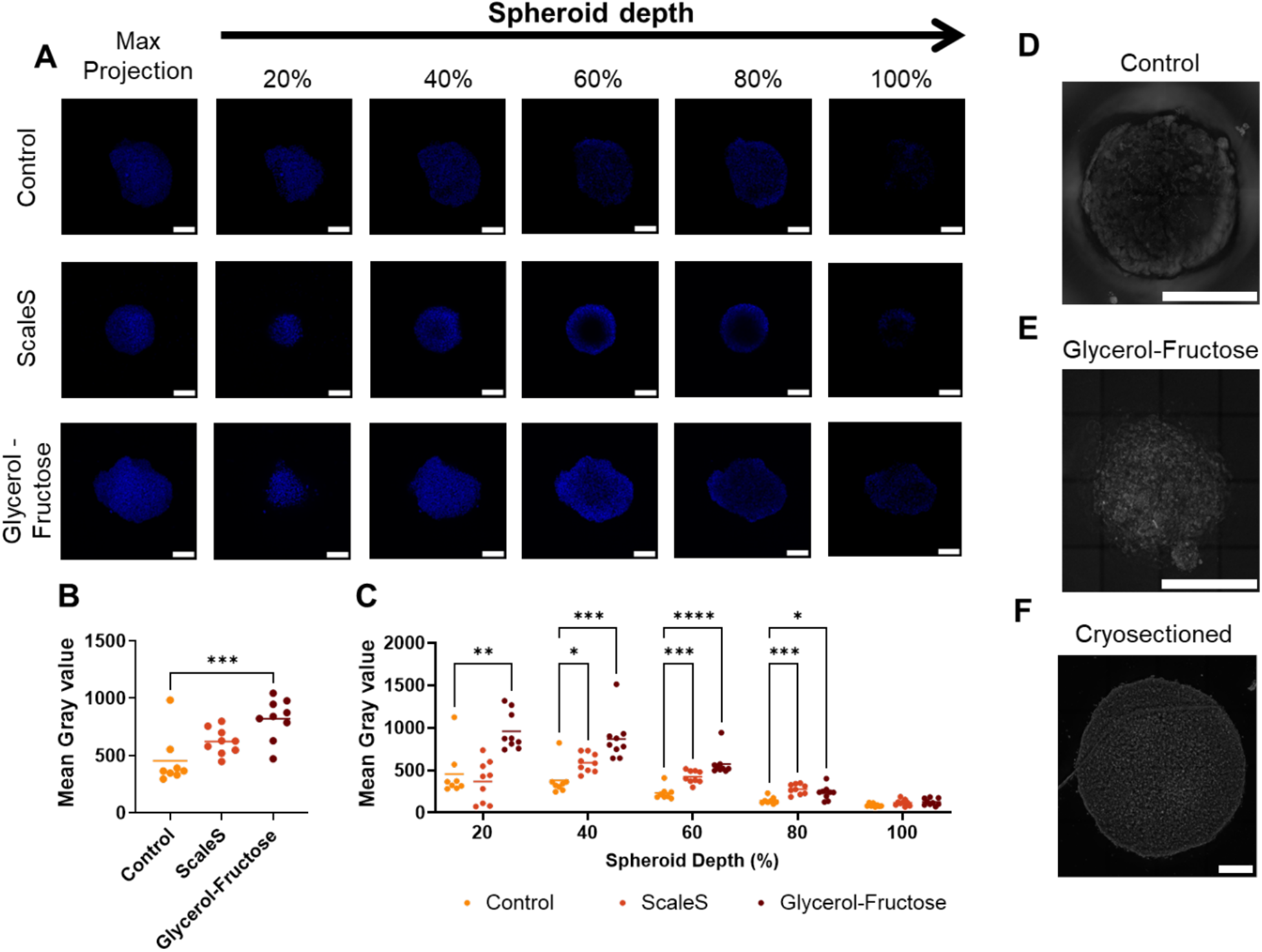
Optical clearing is incompatible with HT analysis. A) Representative fluorescent images of spheroids stained and imaged following different clearing protocols. Images include a maximum-intensity projection and maximum-intensity projections through defined intervals of spheroid depth. B) Quantification of Mean Gray Value measured from maximum-intensity projections, Data are represented as individual spheroids with the mean (n = 2-3 spheroids per condition from three biological replicates). Statistical significance was determined by one-way ANOVA with Šídák’s post-hoc test. C) Quantification of MGV from maximum-intensity projections at different spheroids depths. Statistical significance was determined by two-way ANOVA with Šídák’s post-hoc test. Representative HT images of spheroids D) without clearing and E) following glycerol-fructose clearing. G) Representative HT images from cryosectioned spheroids. Scale bar: 100 µm. * p ≤ 0.05, ** p ≤ 0.01, *** p ≤ 0.001, **** p ≤ 0.0001 vs. control. HT = Holotomography.

The glycerol–fructose protocol was then applied to HT imaging and compared with permeabilised controls (Fig. 1D,E). Despite the positive results observed using confocal microscopic imaging, RI-matching resulted in a reduction of the intrinsic image contrast, which compromised accurate spheroid segmentation by the analysis software (Fig. 1E). We therefore applied HT to spheroids that had been cryosectioned without a clearing agent. These sectioned spheroids yielded clear holographs compatible with quantitative HT analysis (Fig. 1F). All subsequent HT analyses were consequently performed on spheroid sections.

### 3.3 EPDCs Reduce Protein Content in Cardioids

Following optimisation, protein concentration, dry mass, and mean refractive index were quantified across both spheroid models, Cardioids and EpiCardioids, using HT imaging of cryosections (Fig. 2A).

**Figure 2.**
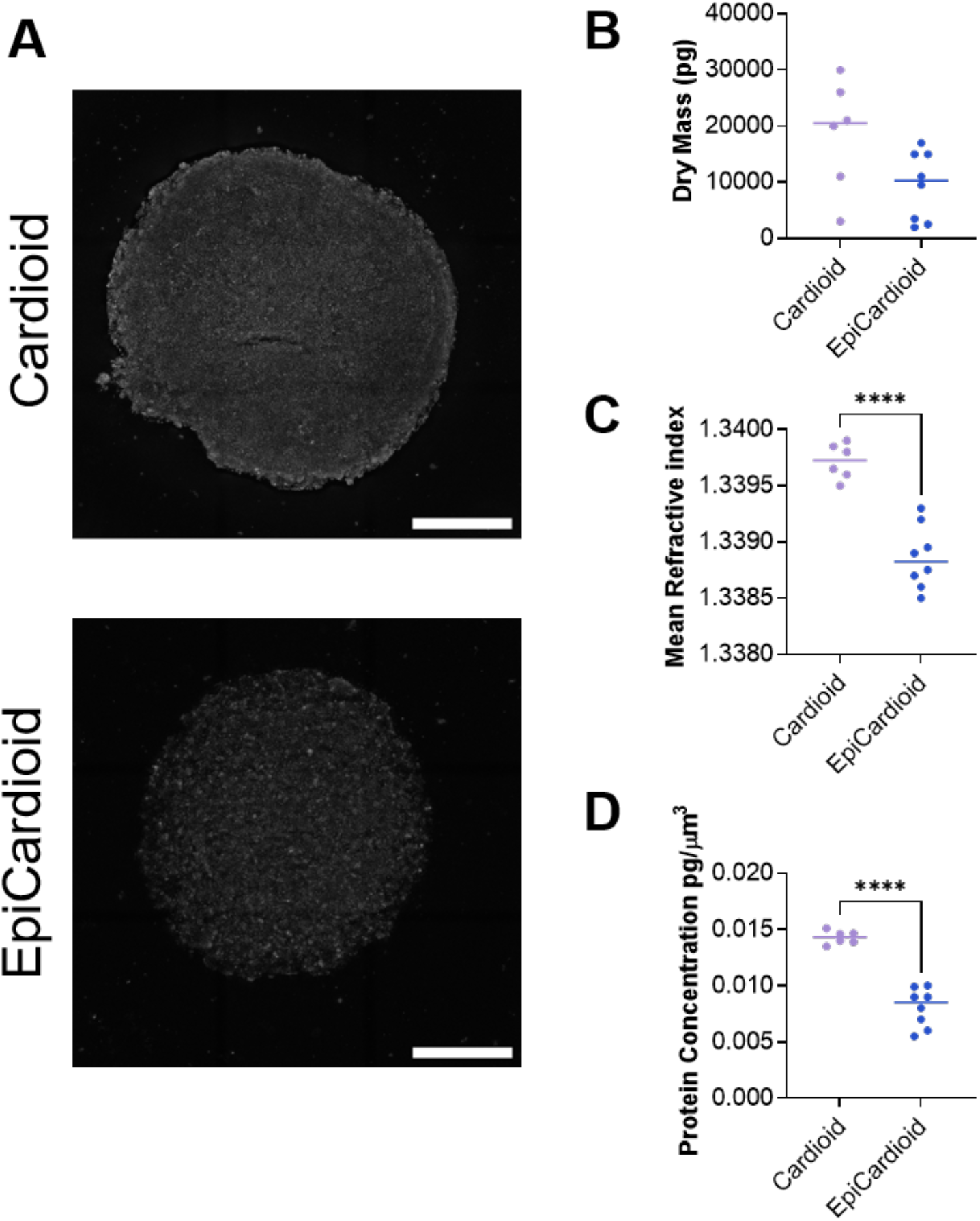
EPDCs reduce protein content in cardioids. A) Representative HT images of cryosectioned spheroids. Analysis of B) dry mass, C) refractive index, and D) protein concentration extracted from spheroids using TomoAnalysis. Scale bar, 100 µm. Data are presented as individual spheroids with the mean (n = 2-3 spheroids per condition from 3 biological replicates). Statistical significance was determined by Students’ t-test. **** p ≤ 0.0001. cardioids = cardiac spheroids, epicardioids = epicardial spheroids.

Dry mass did not differ significantly between Cardioids and EpiCardioids, suggesting that the addition of EPDCs did not significantly alter total biomass (Fig. 2B). However, EpiCardioids exhibited a significant reduction in mean refractive index, corresponding to a significant decrease in intracellular protein concentration relative to Cardioids (Fig. 2C, D). Together, these findings suggest that the incorporation of EPDCs exerts a measurable influence on the biochemical properties of the spheroids.

### 3.4 Hypoxia-Reperfusion Reduces Spheroid Protein Concentration

HT was used to identify changes in protein concentration in spheroids subjected to hypoxic-reperfusion (H/R). Cardioids and EpiCardioids were exposed to hypoxia (<1%) for 16 h followed by 24 h of reperfusion, and were compared by HT imaging with spheroids cultured under normoxia (Fig. 3A, B). HT analysis revealed no changes in overall dry mass (Fig. 3C), however, H/R produced a statistically significant decrease in mean refractive index and the corresponding protein concentration in Cardioids (Fig. 3D, E). For comparison, the bicinchoninic acid (BCA) assay, a standard biochemical method for quantifying total protein, was performed on pooled spheroid lysates. Although visible trends were apparent, none reached significance (Fig. 3F), with absorbance values near the lower detection limit of the assay. In contrast, no significant differences in any measured parameter were observed in EpiCardioids following H/R (Fig. 3B–E).

**Figure 3.**
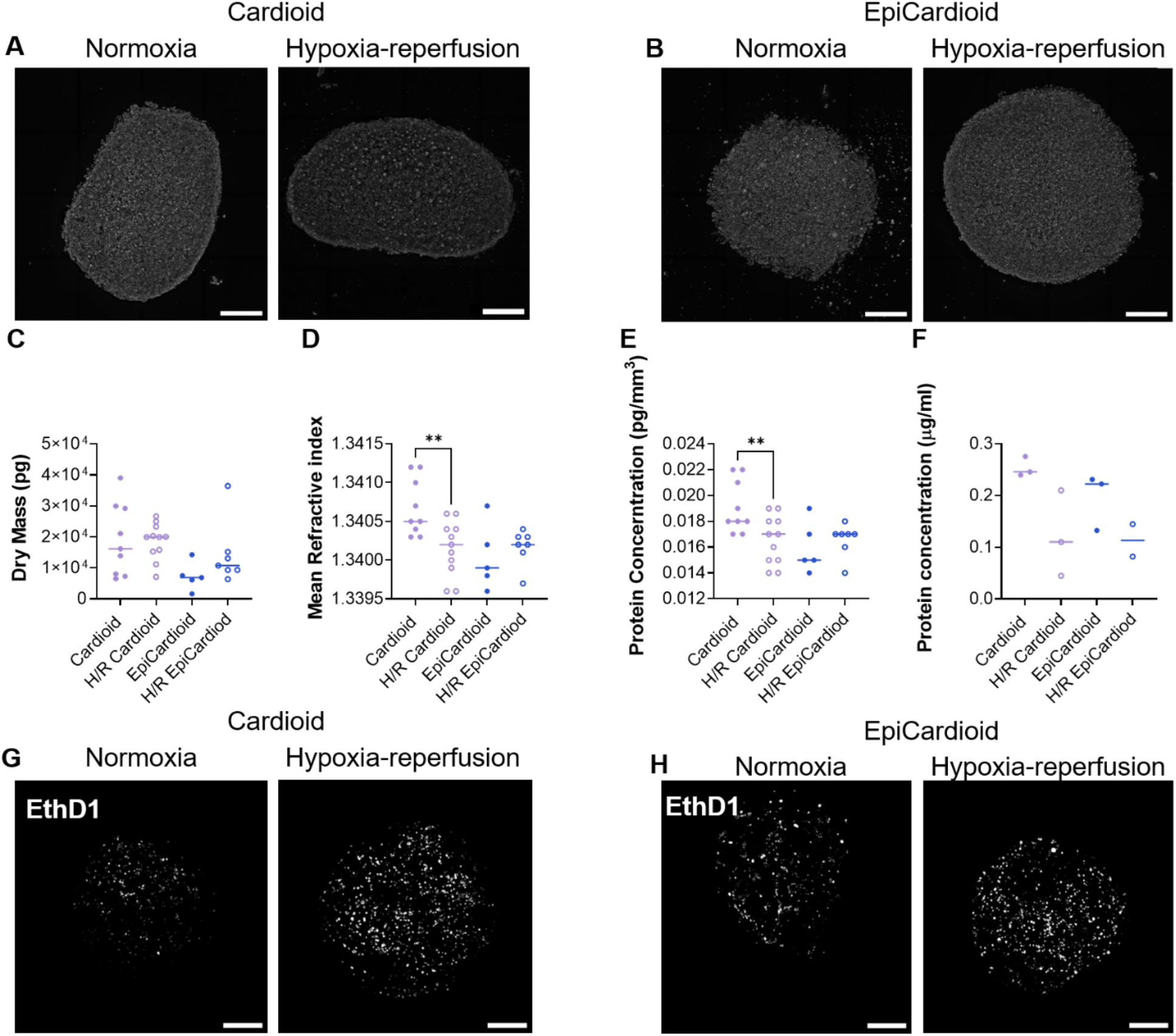
HT detects a reduction in protein content following hypoxia. Representative HT images of cryosectioned A) Cardioids and B) EpiCardioids. Analysis of C) dry mass, D) mean refractive index, and E) protein concentration extracted from spheroids using TomoAnalysis. F) Protein concentration of pooled spheroids obtained from BCA assay. Representative confocal images of G) Cardioids and H) EpiCardioids stained with EthD1. Data are presented as individual spheroids with the mean (n = 2-4 spheroids per condition from 3 biological replicates). Statistical significance was determined by one-way ANOVA of preselected pairs. Scale bar: 100 µm. ** p ≤ 0.01. H/R=hypoxia-reperfusion, cardioids=cardiac spheroids, epicardioids=epicardial spheroids.

To determine whether the changes in protein concentration can be attributed to differences in spheroid viability, spheroids were stained with ethidium homodimer-1 (EthD-1). Representative images showed a reduction in viability following H/R in both Cardioids and EpiCardioids (Fig. 3G, H), indicating that cell death occurred in both models. The preservation of protein concentration in EpiCardioids despite equivalent viability loss therefore suggests that EPDCs contribute a distinct, injury-responsive mechanism that sustains bulk protein content independently of cell survival.

Overall, HT imaging reveals that H/R selectively depletes protein concentration in Cardioids but not EpiCardioids, despite comparable cell death in both models.

## 4. Discussion

Multicellular spheroids are increasingly recognised as powerful *in vitro* models of the cardiac microenvironment, capturing aspects of both developmental biology and disease that are poorly represented in traditional 2D cultures^2,15,26^. In this study, we set out to develop 3D cardiac spheroids that were specifically optimised for HT imaging. This method is quantitative and optimisation of this technique on spheroids would therefore allow innovative biochemical analysis^19^.

To identify the most appropriate formation method, we compared four commonly used spheroid generation techniques: hanging drop, pellet formation, magnetic levitation, and ultra-low attachment (ULA) culture. Hanging drop and pellet methods reliably produced viable spheroids, however, both generated structures with considerable size heterogeneity and frequently yielded multiple spheroids per well. This variability complicates control over initial cell number and composition, directly limiting the interpretability of quantitative imaging readouts. These observations are consistent with previous reports documenting the morphological variability and limited scalability of these approaches, including challenges associated with media evaporation, the difficulty of performing media changes, and shear stress-associated damage during handling^27–29^.

In light of these limitations, we focused subsequent optimisation on magnetic levitation and ULA-based methods, which offered better control over singular spheroid composition and uniformity. Despite these advantages in principle, magnetic levitation proved suboptimal for HT measurements. Although it produced compact and well-defined spheroids, the use of nanoparticle-based levitation reagents caused substantial scattering. The iron-containing NanoShuttle rendered them visibly black and optically opaque which is incompatible with HT as it causes strong absorption and thus, significant scattering, reducing image quality and severely inhibiting quantitative analysis. Although ULA spheroids were not opaque, they still caused significant scattering due to their large size and compact formation ^19,30^

To address these issues, we reduced the spheroids size and applied optical clearing to minimise refractive index mismatch-induced scattering. We evaluated two established aqueous clearing protocols, glycerol-fructose and ScaleS, both of which have been reported to perform comparably to one another and favourably relative to other clearing approaches in fluorescence-based workflows^31^. In our hands, glycerol-fructose yielded a significantly greater improvement in fluorescent signal penetration than ScaleS. Suggesting it is a more effective RI-matching medium in this context. This likely reflects the close correspondence between the RI of glycerol-fructose (1.47–1.53) and that of collagen (1.43–1.53), the predominant structural protein of the cardiac extracellular matrix (ECM), making it particularly well-suited for RI matching in ECM-rich tissue models^31–36^. Although matching the refractive index of the spheroids via optical clearing facilitated light transmission, it also reduced intrinsic contrast in HT reconstructions. Making it more challenging for to segment and quantify. Our results reinforce the idea that clearing protocols must be carefully tailored to the imaging modality and analytical pipeline rather than adopted from fluorescence-based workflows^37^.

Spheroids were sectioned into thinner slices that remained structurally coherent yet significantly less dense and reduced optical path length. This approach allowed us to maintain sufficient RI contrast for reliable segmentation while improving resolution and reducing artefacts associated with out-of-focus scattering. In our hands, sliced spheroids proved superior for HT-based quantification, providing a reasonable compromise between physiological relevance, imaging performance and analysis compatibility. Using this optimised platform, we compared EpiCardioids and Cardioids and observed a significant reduction in mean RI, and by extension protein concentration, in EpiCardioids relative to Cardioids. One mechanistically plausible explanation is that epicardial-derived cells (EPDCs) actively remodel the surrounding ECM through secretion of matrix metalloproteinases (MMPs), a process well-documented in the context of epicardial EMT and cell migration. MMP-mediated matrix degradation would be expected to reduce ECM protein density which would lower the average RI^38,39^. However, since HT provides a bulk RI measurement, it cannot discriminate between contributions from structural proteins, signalling molecules or cell death. Future work combining HT with quantitative proteomics and ECM composition assays in matched EpiCardioid and Cardioid models will therefore be required to define the specific molecular changes underlying this difference.

We then subjected both spheroid types to hypoxia-reperfusion (H/R) to mimic myocardial infarction and reperfusion injury, and used HT to quantify changes in protein concentration. Following H/R, we observed a marked drop in RI and protein concentration in Cardioids, consistent with reduced protein density and increased cell death which is to be expected following hypoxia-reperfusion^40,41^. In contrast, mean RI in EpiCardioids remained largely unchanged after H/R. Critically, EthD-1 viability staining demonstrated a comparable reduction in cell viability in both Cardioids and EpiCardioids following H/R, confirming that the preservation of protein concentration in EpiCardioids is not attributable to differential cell death. This dissociation between viability loss and protein concentration in EpiCardioids suggests that EPDCs contribute an active, injury-responsive mechanism that sustains bulk protein content independently of cell survival. One possible explanation is that EPDCs become reactivated upon injury and secrete a repertoire of paracrine factors, including retinoic acid, hepatocyte growth factor, vascular growth factors, fibroblast growth factors, and transforming growth factors, which support survival, growth and maturation of neighbouring cells^42,43^. The accumulation of these signalling proteins within the spheroid may partially mask the loss of structural proteins from dying cells at the level of bulk RI measurements. An investigation into the protein makeup of these spheroids and the signalling factors present in the supernatant should be assessed to determine how injury-induced paracrine signalling and matrix remodelling shape the bulk protein concentration read out by HT.

To compare HT-derived protein concentration estimates against a conventional method, we performed BCA protein assays on spheroid lysates. Absorbance values were consistently low and approached the limit of reliable detection, likely reflecting the practical difficulties of achieving complete lysis of dense 3D structures. Whilst this precluded a direct quantitative comparison with HT-derived values, it highlighted a fundamental limitation of bulk biochemical assays when applied to small, dense microtissue models and further supports the case for label-free imaging as a complementary or preferable approach for protein quantification in this context.

### 4.1 Limitations

Several limitations must be acknowledged when interpreting these findings. Access to the HT microscope was restricted, limiting the sample size achievable for imaging experiments. Ideally, a minimum of three spheroids per replicate would be required to ensure reproducibility. However, this was not consistently achievable within the available imaging time.

In addition, because spheroids were fixed and cryosectioned prior to imaging, the proteins within the tissue were crosslinked during fixation, meaning that the protein concentration measurements cannot be directly extrapolated to the protein composition of live spheroids. Cryosectioning itself introduced a further source of variability. Due to the inherent limitations of this technique, it was not possible to precisely identify or consistently reproduce the depth of the spheroid being imaged across sections and replicates. Central sections were preferentially selected to minimise variation, as protein concentration and cellular composition are known to vary with spheroid depth because of the oxygen and nutrient gradients that exist within three-dimensional structures^44^.

## 5. Conclusion

In conclusion, by systematically evaluating spheroid generation, optical clearing, and sectioning strategies, this study provides a practical and transferable workflow for HT imaging of cardiac microtissues. We show that HT sensitively detects biologically meaningful differences in protein concentration between spheroid compositions and in response to H/R injury. H/R drove a significant reduction in bulk protein concentration in Cardioids but not in EpiCardioids, suggesting an active role for the epicardial layer. These findings establish a foundation for future work integrating HT-derived biophysical readouts with molecular and proteomic assays to advance understanding of cardiac injury and repair in 3D models.

## Acknowledgements

This work was supported by the UK Engineering and Physical Sciences Research Council (EPSRC) through a Doctoral Training Partnership award (grant number EP/W524463/1). We also thank Tomocube Inc., the company that commercialised the HT system used in this study, for valuable scientific discussion and expert advice. Additionally, H9C2 cells were generously provided by Dr Ioannis Smyrnias at the University of Surrey.

## Resource availability

Lead contact Material availability Data availability

## Declaration of generative AI and AI-assisted technologies in the writing process

During the preparation of this work, the author used perplexity in order to improve flow and grammar. After using this tool the author reviewed and edited the content as needed and takes full responsibility for the content of the publication.

